# IHRW: An Improved Hypergraph Random Walk Model for Predicting Three-Drug Therapy

**DOI:** 10.1101/2021.02.25.432979

**Authors:** Qi Wang, Guiying Yan

## Abstract

Drug combination therapy is a well-established concept in the treatment of complex diseases due to its fewer side effects, lower toxicity, and better efficacy. However, it is challenging to identify efficacious drug combinations from many drug candidates. Computational models could greatly reduce the cost, but most models did not use data for more than two-drug combinations and could not predict three-drug therapy. However, three-drug combinations account for about 21% of the known combinations, which is a very important type of treatment. Here, we utilized higher-order information and developed an improved hypergraph random walk model (IHRW) for three-drug therapy prediction. This is the first method to explore the combination of three drugs.

As a result, the case studies of breast cancer, lung cancer, and colon cancer showed that IHRW had a powerful ability to predict potential efficacious three-drug combinations, which provides new prospects for complex disease treatment. The code of IHRW is freely available at https://github.com/wangqi27/IHRW.

## Introduction

Drug combinations have been widely used in the treatment of complex diseases, such as cancer and AIDS(Chou, 2006). There is a growing enthusiasm for the development of potentiated drug combinations, both in academic institutions and in the pharmaceutical industry. For example, CRx-102, which has completed phase II clinical trials, is a combination of dipyridamole and low-dose prednisolone, which can be used to treat osteoarthritis (Zhang *et al*., 2009). The underlying reason for the enthusiasm for combination studies is that potentiating drug combinations can increase the therapeutic efficacy at the same dosage and decrease the dosage at the same efficacy, thus reducing drug toxicity. Of course, drug combinations have other advantages that single drugs do not have, such as simultaneous modulation of multiple targets, populations, and diseases (Zimmermann *et al*., 2007) and delaying the development of resistance (Chou, 2006)(Zimmermann *et al*., 2007). These advantages have led to an increasing number of researchers investigating drug combinations (Borisy *et al*., 2003)(Zimmermann *et al*., 2007). From the above discussion, it can be concluded that the prediction of potent drug combinations is an important issue for the improvement of human health care.

If there are a total of *n* drug candidates, then there are 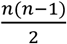 pairs of combinations consisting of two drugs, and many higher-order combinations (Borisy *et al*., 2003). It is conceivable that even a very small number of drug candidates will lead to many combinations (Borisy *et al*., 2003). Therefore, trying all possible combinations would pose a huge challenge in terms of time and cost for ornamentation and thus would not be feasible. Even with high-throughput screening techniques, limited drug combination experiments can only sample a small fraction of the total drug combination space (Zhang *et al*., 2007). Therefore, it is difficult to find the optimal drug combination or the optimal drug combination pair concentration. The need for predicting potential potent drug combinations based on database and literature information is very urgent. Computational prediction methods can reduce the number of trials and thus the time and cost of experiments. We only need to perform rigorous biological tests on the predicted potentially potent drug combinations. In this sense, prediction methods guide drug combination experiments and facilitate the understanding of drug combination mechanisms.

Recently, several computational approaches have been proposed for the prediction of efficacious drug combinations. Bai et al. (Bai *et al*., 2018) used an improved Naïve Bayesian algorithm to predict effective drug combinations. Huang et al. (Huang *et al*., 2014) developed a network-based ranking algorithm DrugComboRanker to construct a disease signaling network using patient genomic information and protein interaction data and to model drug functional networks based on drug genomic profiles. Chen *et al*. (Chen *et al*., 2016) developed a model named Network-based Laplacian regularized Least Square Synergistic drug combination prediction (NLLSS) to predict potential synergistic drug combinations. They also conducted biological experiments, and finally, 7 out of 13 antifungal synergistic drug combinations predicted by NLLSS were validated.

However, most computational models did not use the data for more than two-drug (i.e., three-drug and four-drug) combinations and could not predict three-drug therapy. The combination of the three drugs has fewer complications and lower mortality than the combination of multiple drugs. This retrospective study (Gaziev *et al*., 2001) showed that the combination of three or more drugs is safe and effective for the treatment of moderate or severe cGVHD at least in younger patients. Therefore, we propose a method called IHRW to combine known potent drug combinations and unlabeled drug combinations for large-scale prediction of potential efficacious drug combinations. Applying IHRW to cancer-related drug combination prediction, the algorithm achieves good results in case studies. This is a real work of predicting potent drug combinations purely based on database and literature information, without relying on experiments.

## Materials and methods

### 2.1 Data collection and pre-processing

The Drug Combination Database (DCDB) (Liu *et al*., 2014)was the first specialized database to collect and organize information on drug combinations, which contains 1363 drug combinations consisting of 1735 drugs. The drug combinations corresponding to breast, lung, and colon cancers in DCDB are the dataset studied in this paper. In detail, the data in the database were selected according to the following criteria. ICD10 is an internationally harmonized disease classification developed by WHO (Quan *et al*., 2005), and each drug combination in the DCDB database has at least one ICD10 code. If a drug combination has an ICD10 code of C18, i.e., colon malignancy, we record this data as lung cancer data. Similarly, drug combinations with ICD10 codes C50 and C34 were recorded as breast cancer and lung cancer data, respectively. We did not include drug combinations with an effective type of “ Need further study” in the DCDB because the experimental results of these drug combinations are unclear.

According to the data in the DCDB database, the drug combination of three-drug accounts for about 22%. This indicates that the existing 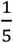 drug combinations cannot be predicted by the previous mathematical model. **Table 1** shows the number and proportion of different single drugs in the combinatorial drugs from the DCDB database.

**Table.1.** The number and proportion of different single drugs of all combinations.

|  | 2-drug | 3-drug | 4-drug | 5-drug | 6-drug | Total |
| --- | --- | --- | --- | --- | --- | --- |
| # of combinations | 946 | 295 | 91 | 25 | 6 | 1363 |
| Percentage | 69.4% | 21.6% | 6.7% | 1.8% | 0.4% | 100% |

### 2.2 Notations and definitions

A hypergraph is a generalization of a graph where an edge could have more than two vertices. By modeling the drug combination data as a hypergraph, we capture the relationship between drugs where we set drugs as vertices and the combination relationships as hyperedges.

Let *HG*(*V, E*) be a hypergraph with the vertex set *V* and the hyperedges set *E. E* is a subset of *V* where ∪_*e*∈*E*_ *e*= *V*. A hyperedge *e* is said to be incident with *v* when *v* ∈ *e*. A hypergraph has an incidence matrix 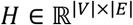 as follows:

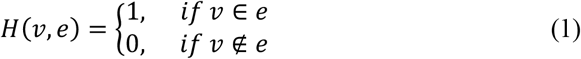

The vertex degree and hyperedge degree are defined as follows:

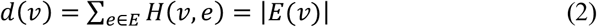

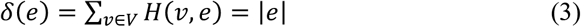

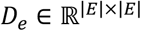 and 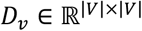 are diagonal matrices that represent the degree of the hyperedges and vertices, respectively.

### 2.3 Hypergraph random walk with different restart strategy

The random walk on the hypergraph is a generalization of it on the simple graph. We regard the vertex set {*v*_1_, *v*_2_, …,*v*_n_} as a set of states {*s*_1_, *s*_2_, …,*s_n_*} in a finite Markov chain 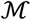. The transition probability of 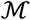 is a conditional probability defined as *P*(*u,v*)=*Prob*(*s*_*t*+1_=*v|s_t_*=*u*) which means that the 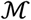 will be at *v* at time *t* + 1 given that it was at *u* at time *t*. What is more, for any *u* of *V* we have ∑_*v*∈*v*_ *P*(*u,v*)= 1.

Define the transition probability 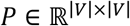 as follows:

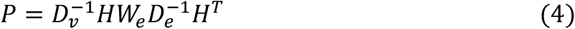

Define 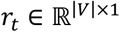 as a vector in which the *i*-th element represents the probability of discovering the random walk at vertex *i*. at step t, so the probability *r*_*t*+1_ can be calculated iteratively by:

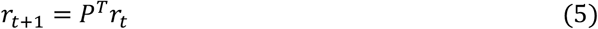

To represent the relationship between multiple nodes, we introduce a restart item. Compared with the above algorithm, there is an additional restart item. The probability *r*_*t*+1_ can be calculated iteratively as follows:

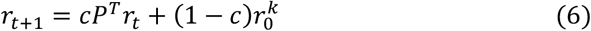

Define initial probability 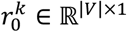 as a vector with the *i*-th element is equal to 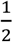 and *k*-th element is equal to 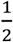, while other elements are zeros. 1 − *c* is the restart probability (0 ≤*c*≤ 1). *c* is the hyperparameter, and taking a similar approach to the previous study (Wang and Yan, 2020), we set *c* to 0.8.

In this paper, the random walk on a hypergraph is a transition process of the hypergraph as shown in figure 1. The IHRW algorithm is convergent (see Section 2.5 for proof), so we can calculate the steady-state probability. When the random walk reaches a steady-state, steady-state probability between vertex *i*. and vertex *j* is defined as the *j*-th element of *r*_t_ corresponding to the starting vertex is *v_i_*.

**Fig. 1.**
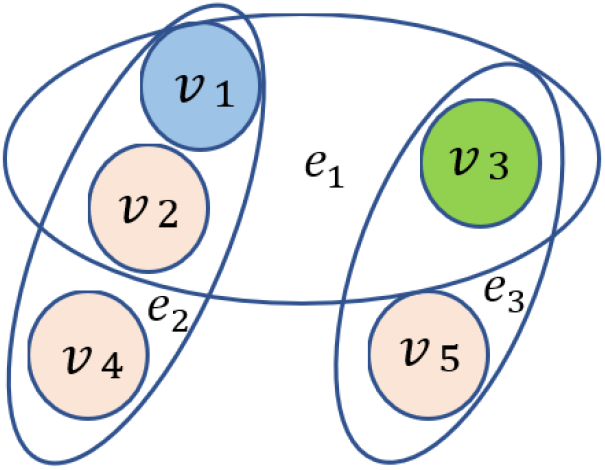
An example of the IHRW. Suppose we want to score a combination of triples (*v_i_, v_j_, v_k_*), the restart probability is 1 − *c*. First, we need to assume two fixed vertices *v_i_* (initial vertex) and *v_k_* (preset vertex), such as *v_i_* = *v*_1_ and *v_k_*= *v*_3_. Then, the initial vertex is *v*_1_, it has *c*/2 probability to choose *e*_1_ or *c*/2 probability to choose *e*_2_, suppose *e*_2_ is selected, *v*_1_ has 1/3 probability to choose *v*_1_, *v*_2_, or *v*_4_ as the next vertex of the transition process; at the same time, the initial vertex *v*_1_ also has (1 − *c*)/2 probability to choose *v*_1_ and (1 − *c*)/2 probability to choose *v*_3_ as the next vertex of the transition process. Especially, if the initial node and the preset node are the same e.g., *v*_1_, then *v*_1_has a 1 − *c* probability of transferring to *v*_1_. Next, repeat the process of “ select hyperedge then select vertex or restart to initial vertex /preset vertex”. Finally, when the number of random walks is enough (the number of the above process tends to infinity), we can obtain the “selected” probability of each vertex, that is, the steady-state probability. For example, when *v_i_* = *v*_1_ and *v_k_*= *v*_3_, *v_i_* = *v*_1_ and *v_k_*= *v*_2_, or *v_i_* = *v*_2_ and *v_k_*= *v*_3_, the steady-state probability of *v*_2_, *v*_3_, and *v*_1_ are 0.3, 0.2, and 0.5 means that the score of triples (*v*_1_, *v*_2_, *v*_3_) is (0.3 + 0.2 + 0.5)/2 = 0.5.

### 2.4 Efficacious scoring of three-drug combinations

Each three-drug combination can be ranked twice according to IHRW. Define Π={π_ijk_}_|*v*|×|*v*|×|*v*|_ be the steady-state probability matrix where *π_ijk_* indicates the steady-state probability among vertex *v_i_*, *V_j_*, and *v_k_*, that is, IHRW starts from vertex *v_i_*, the preset vertex is *v_k_*, and the probability of reaching vertex *v_j_* when the process reaches steady-state. Briefly speaking, the value of *π_ijk_* is the *j*-th element of steady-state probability 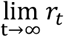 when the *i*-th and *k*-th element of *r*_0_ are both 1/2. The significance of *π_ijk_* can be understood as the possibility of the efficacious drug combination of drug *j* (i.e., for *n* − 2 candidate drugs except for drug *i*. and *k*) with given drug i and drug *k*. The goal of IHRW is to predict the efficacious combination of any three drugs, so each drug cannot be combined with itself, but can be combined with other n − 1 drugs. Therefore, the efficacious score *S*={*S_ijk_*}_|*v*|×|*v*|×|*v*|_ is defined as follows:

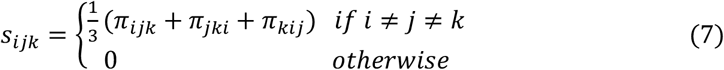

The efficacious score of any three drugs can be calculated by IHRW, and it can be expected that the three-drug combination with the higher efficacious score has a higher probability of becoming a combinatorial drug. Experimenters can prioritize the validation of these three-drug combinations, and therefore the cost of identifying potential drug combinations can be significantly reduced. The flowchart of the IHRW is shown in **Figure 2**.

**Fig. 2.**
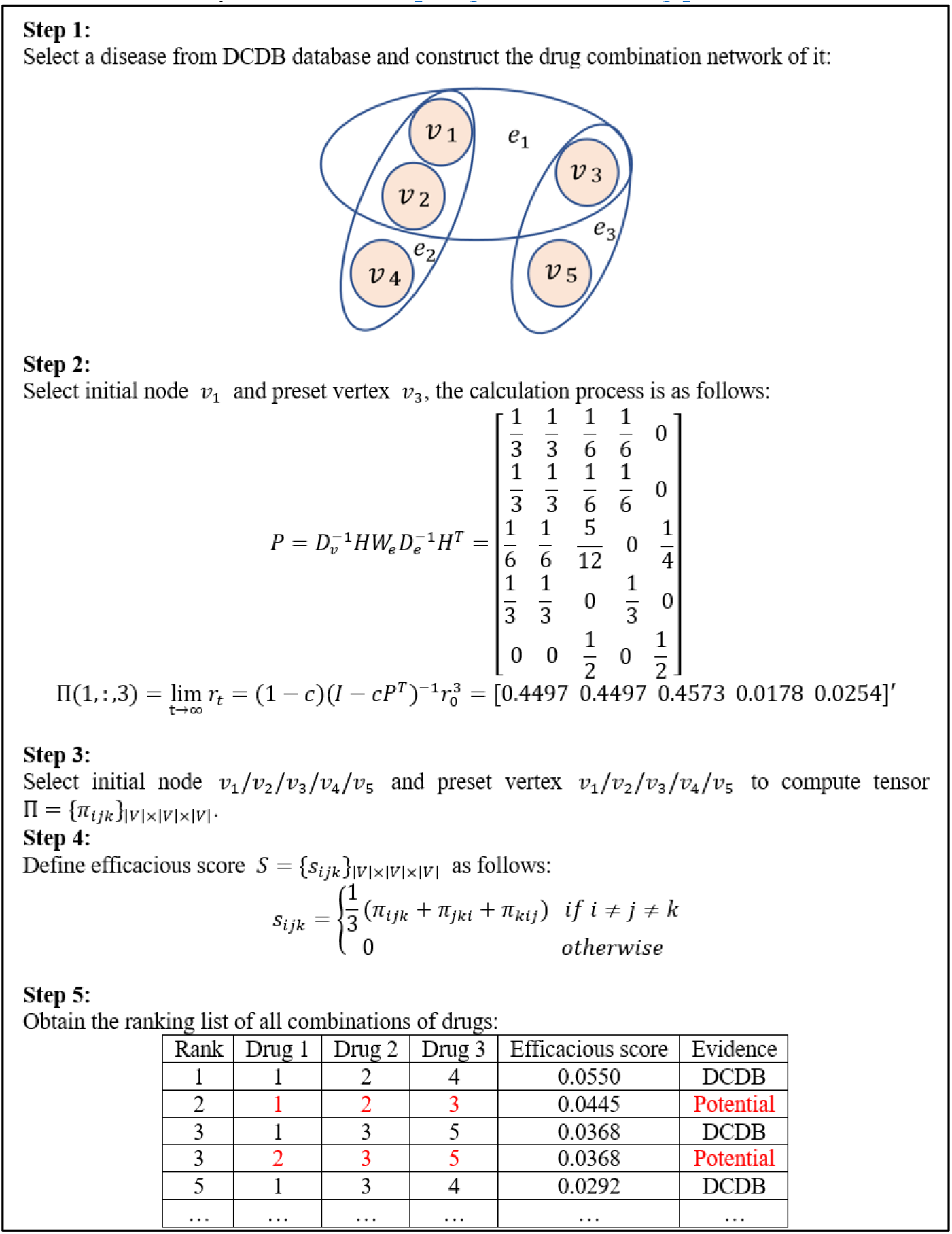
Flowchart of the IHRW.

### 2.5 Convergence of the algorithm

Here we will prove that the IHRW algorithm is convergent, that is 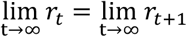.

For any t ≥ 0, We first have

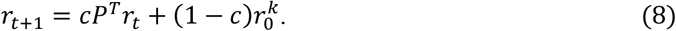

Define

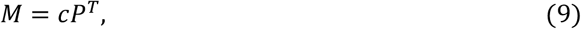

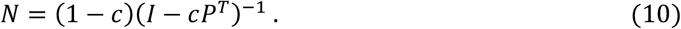

Thus,

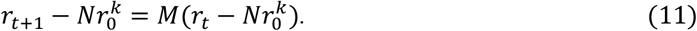

Then, define

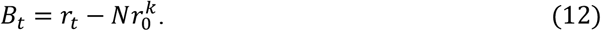

Therefore, (11) can be written as follows:

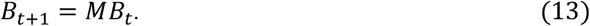

By (12), when t = 0, we have 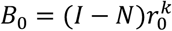, thus

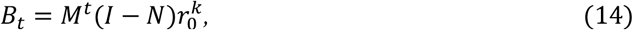

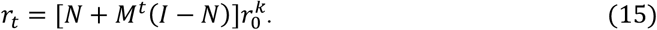

Since 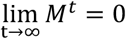, we have

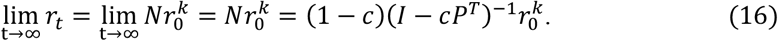

Hence,

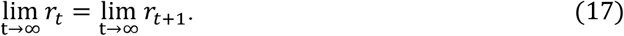

## Results

### 3.1 Basic network characteristics

The characteristics of the drug combination network are shown in **Table 2**. Here, hyperedge represents a known combination of efficacious drugs and vertex represents the drugs involved in the combination of efficacious drugs. The number of hyperedges with three vertices represented the number of the efficacious drug combination of three drugs.

**Table.2.**
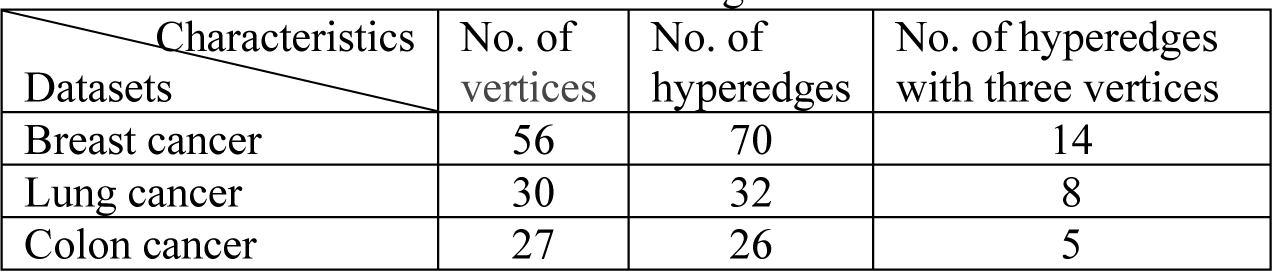
Characteristics of the drug combination network.

| Characteristics<br>Datasets | No. of<br>vertices | No. of<br>hyperedges | No. of hyperedges<br>with three vertices |
| --- | --- | --- | --- |
| Breast cancer | 56 | 70 | 14 |
| Lung cancer | 30 | 32 | 8 |
| Colon cancer | 27 | 26 | 5 |

### 3.2 Analysis of the reasonableness of scoring

We compared the mean scores of known three-drug combinations with the mean scores of all possible combinations (**Table 3**), and the result was that IHRW scored known combinations higher than the mean scores of all possible combinations, indicating that IHRW has some predictive power and can clearly distinguish currently known drug combinations from uncertain ones.

**Table.3.**
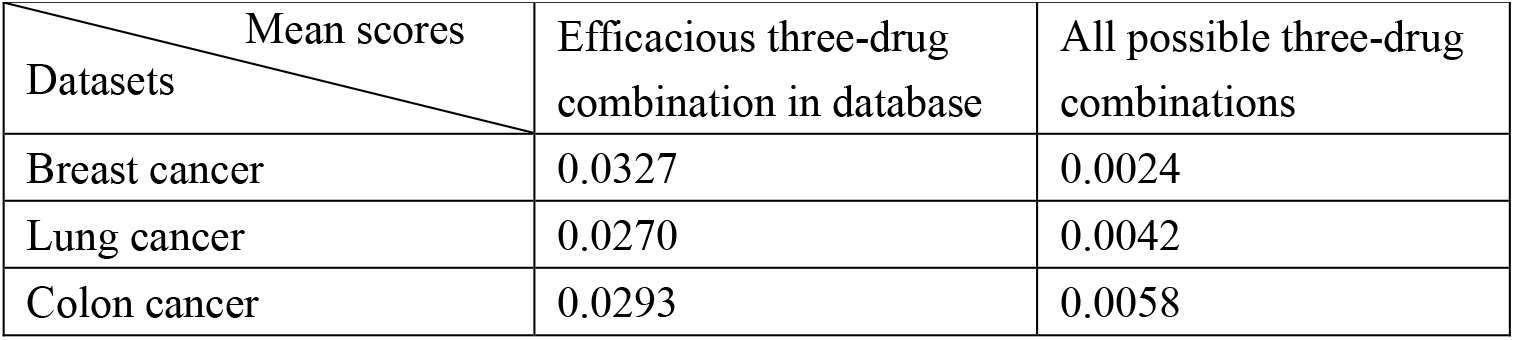
Comparison of the average scores of known and all possible three-drug combinations.

**Table.4.** Case study of breast cancer

| Rank | Drug1 | Drug2 | Drug3 | Evidence |
| --- | --- | --- | --- | --- |
| 1 | Palonosetron | Dexamethasone | Aprepitant | DCDB |
| 1 | Dexamethasone | Granisetron | Prochlorperazine | DCDB |
| 3 | Vitamin D | Calcium | Zoledronate | DCDB |
| 4 | PD98059 | Gefitinib | Taxane | Null |
| 5 | Raloxifene | Fluorouracil | Trimetrexate | DCDB |

**Table.5.**
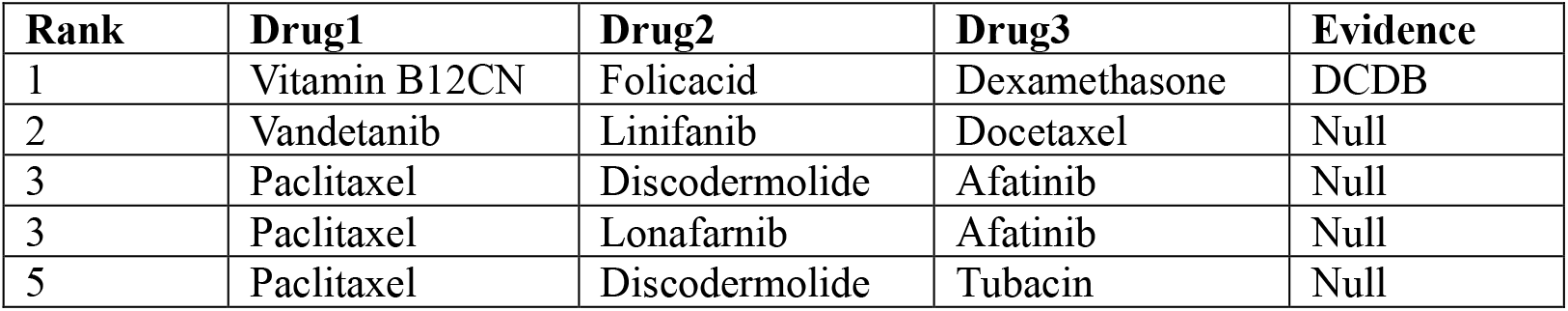
Case study of lung cancer

### 3.3 Case studies

In this section, based on the gold standard dataset, we conducted case studies to validate the efficiency of IHRW in discovering potential combination drugs. For breast cancer, lung cancer, and colon cancer, we ranked all possible three-drug combinations based on their corresponding predicted efficacious scores. The predictions were validated not only against the DCDB database but also against recently published experimental literature. **Table 2** shows that efficacious three-drug combinations represent approximately a very small proportion of the total combinations in each cancer dataset, therefore, we analyzed the top 5 predictions. The “Null” in the table indicates that documentation has not been found.

Breast cancer is the most common cancer and the leading cause of cancer-related deaths in women worldwide(Zheng *et al*., 2014). 14 groups of efficacious three-drug combinations have been discovered by biological experiments, IHRW can also predict more potential efficacious three-drug combinations. Four of the top five scoring drug combinations were validated as efficacious. The combination of 100 mg/kg Gefitinib and 20 mg/kg Taxane showed a strong synergistic effect on MCF7/ADR cells, but an invitro additive-antagonistic effect on MDA-MB-231 cells. Gefitinib-Taxane is better than monotherapy. The literature (Takabatake *et al*., 2007) suggests that the possible mechanism of Gefitinib-Taxane is that Taxane produced the anticancer effect by inducing apoptosis and microtubule disruption. Crosstalk between EGFR and hypoxia-inducible factor-1alpha pathways increased resistance to apoptosis by upregulating survivin. Gefitinib produced an anticancer effect via EFFR tyrosine kinase inhibition, which offsets the counteractive EGFR-hypoxiacrosstalk in resisting taxane’s pro-apoptosis activity. What is more, (Normanno *et al*., 2006) showed that the combination of PD98059 and Gefitinib was efficacious to breast cancer. According to the literature, PD98059 (MEK1 inhibitor) has been shown to act as a highly selective inhibitor of MEK1 activation and MAP kinase cascade in vivo. PD98059 binds to the inactive form of MEK1, preventing the activation of upstream activators such as c-Raf. MEK1 and MEK2, also known as MAPK or Erk kinases, are bispecific protein kinases that function in a mitogen-activated protein kinase cascade to control cell growth and differentiation. Although the combination of PD98059-Gefitinib-Taxane has not yet been experimentally validated as effective against breast cancer, we believe that the effectiveness of this combination may be verified shortly.

Lung cancer begins with a mutation of normal cells that turns them into cancer cells. These cells divide and multiply at an uncontrolled rate and eventually form tumors that block breathing and oxygen supply to the entire body(Dela Cruz *et al*., 2011). We used IHRW to predict potentially efficacious three-drug combinations. As a result, only one lung cancer-related top-ranked efficacious triple-drug combination was validated by the database and recent experimental literature. But literature (Herbst *et al*., 2010; Grève *et al*., 2012; Honore *et al*., 2004) showed that the combination of Vandetanib-Docetaxel, Paclitaxel-Afatinib, and Paclitaxel-Discodermolide are efficacious to non-small-cell lung cancer. (Shi *et al*., 2000) showed that the combination of Paclitaxel and Lonafarnib is efficacious to human lung tumor and mammary tumor. (Marcus *et al*., 2005) showed that Paclitaxel-Tubacin is efficacious to lung and breast cancer. The 2-drug combination between them is efficacious in lung cancer, and we believe that the 3-drug combination between them is likely to be efficacious as well.

Colon cancer is one of the most common malignant tumors in the world (Xue *et al*., 2015), killing almost seven hundred thousand people every year (Gu *et al*., 2017). From **Table 6**, we could know that only one efficacious three-drug combination in the data set had been predicted by IHRW. First of all, the combination of Tegafur, 5-2-4-dihydroxypyrimidine, and Potassium oxonate ranked top one by IHRW but it is not validated by databases and recent experimental literature. (Shirasaka *et al*., 1996) evaluate practically the antitumor activity of 1 M tegafur-0.4 M 5-chloro-2,4-dihydroxypyridine-1 M potassium oxonate (S-1), a new p.o. fluoropyrimidine, in comparison to 1 M tegafur-4 M uracil, the findings suggested that this nude mouse orthotopic human colorectal tumor model is useful for evaluating the clinical efficacy of drugs or therapies for colorectal cancer and that S-1 is more effective than UFT in treating human colorectal tumors. Since this drug combination is effective in nude rats, we are confident enough to believe that it can be validated for humans in the future. Secondly, the combination of RPR-115135, 2’-Deoxyinosine, and Fluorouracil is not recorded in the database of DCDB, but the combination of RPR-115135 and Fluorouracil is recorded. (Russo *et al*., 2002) showed that RPR-115135 significantly enhances the efficacy of 5-FU only when p53 is functioning. The possible mechanism is that joint tumor suppressive (via fluorouracil stabilization of P53) and antiproliferative (via RPR-115135 inhibition of Ras farnesylation) actions. Moreover, the combination of Capecitabine, Floxuridine, Dexamethasone, and Oxaliplatin is also recorded in the database of DCDB(Alberts *et al*., 2010). Oxaliplatin binds preferentially to the guanine and cytosine moieties of DNA, leading to cross-linking of DNA, thus inhibiting DNA synthesis and transcription. Capecitabine is a prodrug that is selectively tumor-activated to its cytotoxic moiety, fluorouracil, by thymidine phosphorylase, and thymidylate is the necessary precursor of thymidine triphosphate, which is essential for the synthesis of DNA, therefore a deficiency of this compound can inhibit cell division. Dexamethasone is a glucocorticoid agonist. In addition to binding to specific nuclear steroid receptors, dexamethasone also interferes with NF-kB activation and apoptotic pathways. Floxuridine inhibits thymidylate synthetase, resulting in disruption of DNA synthesis and cytotoxicity. Therefore, we are confident that Capecitabine-Floxuridine-Dexamethasone and Oxaliplatin-Floxuridine-Dexamethasone, both three-drug combinations predicted by the IHRW, are likely to be effective against colon cancer.

**Table.6.** Case study of colon cancer

| Rank | Drug1 | Drug2 | Drug3 | Evidence |
| --- | --- | --- | --- | --- |
| 1 | Tegafur | 5-2-4-dihydroxypyrimidine | Potassium oxonate | Null |
| 2 | Fluorouracil | Methotrexate | Leucovorin | DCDB |
| 3 | RPR-115135 | 2'-Deoxyinosine | Fluorouracil | Null |
| 4 | Capecitabine | Floxuridine | Dexamethasone | Null |
| 5 | Oxaliplatin | Floxuridine | Dexamethasone | Null |

As we can see from **Table 2**, the number of single drugs, drug combinations, and three-drug combinations corresponding to breast cancer is relatively high, resulting in the highest percentage of validated three-drug combinations with higher IHRW scores, so there is a high probability that the drug combinations predicted by lung cancer and colon cancer can be applied to the treatment of cancer.

## Discussion

Most current computational models do not use data from more than two-drug combinations and do not predict three-drug therapy. However, three-drug combinations account for about 21% of the known. As far as the authors know, this is the first model to predict the efficacious combination of more than two drugs, and the combination information of more than two drugs is used in the modeling process, which is more comprehensive.

The IHRW proposed in this paper is a general framework, and its convergence is proved theoretically. For the problem of predicting whether there are three vertices closely related in the graph, the framework can calculate the correlation score for any three vertices. This algorithm has a wide range of application scenarios, such as the research on the relationship between the signal strength of multiple communication base stations, the relationship between multiple proteins, the relationship between different brain regions, and so on.

There are still aspects of the model that could be improved. The choice of restart probabilities did not incorporate the biological context of the drug combinations. The IHRW can predict which drugs can be the components of a three-drug combination, but the dosage of a single drug cannot be predicted, which is an important problem to be overcome in the future. For link prediction of binary relationships, there are common metrics such as Area Under the Receiver Operating Characteristic curve (AUROC), and Area Under the Precision-Recall curve (AUPR) to measure the strength of the algorithm’s prediction ability, but currently, there are no evaluation metrics for link prediction of multivariate relationships. It is hoped that future researchers can develop evaluation metrics for hyperedge link prediction between multiple entities.

